# CLE11 and CLE10 Suppress Mycorrhizal Colonisation in Tomato

**DOI:** 10.1101/2023.02.21.529440

**Authors:** Kate Wulf, Chenglei Wang, Tania Ho-Plagaro, Choon-Tak Kwon, Karen Velandia, Alejandro Correa-Lozano, María Isabel Tamayo-Navarrete, Jiacan Sun, James B. Reid, Jose Manuel García Garrido, Eloise Foo

## Abstract

Symbioses with beneficial microbes are widespread in plants, but these relationships must balance the energy invested by the plants with the nutrients acquired. Symbiosis with arbuscular mycorrhizal (AM) fungi occurs throughout land plants but our understanding of the genes and signals that regulate colonisation levels is limited. Here, we demonstrate that in tomato two CLV3/EMBRYO-SURROUNDING REGION (CLE) peptides, *Sl*CLE10 and *Sl*CLE11, act to suppress AM colonisation of roots. Mutant studies and overexpression via hairy transformation indicate *SlCLE11* acts locally in the root to limit AM colonisation. Indeed, *SlCLE11* expression is strongly induced in AM colonised roots but *SlCLE11* is not required for phosphate suppression of AM colonisation. *Sl*CLE11 may act through as yet uncharacterised signalling pathways, as *SlCLE11* does not suppress AM colonisation by acting through two previously characterised receptors with roles in regulating AM colonisation, *Sl*FAB (CLAVATA1 orthologue) or *Sl*CLV2. *Sl*CLE10 appears to play a more minor or redundant role, as *cle10* mutants did not influence AM, although the fact that ectopic overexpression of *SlCLE10* did suppress colonisation suggests *SlCLE10* may play a role in regulating AM colonisation. Our findings show that CLE peptides regulate AM colonisation in the non-legume species tomato.

## Introduction

Many plants form intimate symbiotic relationships with soil microbes that enhance nutrient uptake. The majority of land plants form a symbiosis with arbuscular mycorrhizal (AM) fungi, and this ancient association is thought to provide the genetic toolkit for the more recently evolved nodulation, the symbioses with nitrogen-fixing bacteria formed by many plants in the fabid clade (reviewed by Radhakrishnan et al. 2020; Wang et al. 2022). The hosting of these microbes is energetically expensive to the plant and this investment must be balanced by benefit gained. Plants monitor the extent of symbioses and use a root-shoot feedback system to limit the extent of microbial colonisation, termed autoregulation of nodulation and autoregulation of mycorrhizae (Chaulagain and Frugoli 2021; Wang et al. 2018). The major macronutrients nitrogen and phosphorus also influence the degree to which plants become colonised (e.g. Isidra-Arellano et al. 2020; Pervent et al. 2021; Wang et al. 2021a).

CLV3/EMBRYO-SURROUNDING REGION (CLE) peptides are short, secreted peptides with important roles in many aspects of plant development and environmental response, including organization of the shoot apical meristem, fruit size, root development and nutrient response (Song et al. 2021; Willoughby and Nimchuk 2021). Studies across a range of legumes have established a major role for a conserved CLE peptide signalling pathway in the control of nodule number (reviewed by Chaulagain and Frugoli 2021; Roy and Muller 2022),. Briefly, nodulation induces the expression of a specific set of CLE peptides and activation of some of these CLE peptides requires arabinosylation via hydroxyproline O-arabinosyltransferase (HPAT) proteins such as *Mt*RDN1 and LjPLENTY. These CLE peptides are perceived in the shoot by potential receptors including leucine-rich repeat receptor-like kinases (LRR-RLK) CLAVATA1 (*Mt*SUNN, *Lj*HAR1, *Ps*SYM29 and *Gm*NARK), *Lj*KLV, *Lj*CRN and a LRR receptor-like protein without a kinase domain, CLV2. This perception activates downstream pathways to suppress further nodulation in the root, including potential roles for shoot-derived cytokinin and suppression of miR2111 which then depresses Too Much Love (TML), an F-Box protein acting in the root. This pathway overlaps at least in part with nitrogen and phosphorous-regulation of nodule number, although independent pathways may also operate (e.g. Isidra-Arellano et al. 2021; Pervent et al. 2021; Roy and Muller 2022).

Studies in legumes reveal conserved genes operate in autoregulation of nodulation and mycorrhizal colonisation, indicating a shared evolutionary history. Common signalling is revealed in split root studies; nodulation can limit subsequent mycorrhizal colonisation and vice-versa (Catford 2003). In addition, although mycorrhizal-specific CLE peptides appear to control mycorrhizal colonisation in legumes, they may act through a common signalling pathways with nodulation. For example, legume mutants disrupted in the CLV1, CLV2 and HPAT proteins outlined above display both supernodulation (reviewed by Roy and Muller 2022), and hypermycorrhizal phenotypes (Karlo et al. 2020; Morandi et al. 2000; Müller et al. 2019; Sakamoto and Nohara 2009; Solaiman et al. 2000). In addition, in *Medicago truncutula, Mt*CLE33 and *Mt*CLE53 function to suppress mycorrhizal colonisation and require both the arabinosylation enzyme *Mt*RDN1 and the CLV1 orthologue *Mt*SUNN (Karlo et al. 2020; Müller et al. 2019). To date, no CLE peptides that regulate mycorrhizal colonisation in non-legumes have been identified. However, mutant studies have demonstrated orthologues of the CLV1 and CLV2 receptors and HPAT enzyme negatively regulate mycorrhizal colonisation rates in tomato (Wang et al. 2021a) and *Brachypodium distachyon* (BdFON1; Müller et al. 2019).

Phosphorous is a major regulator of mycorrhizal colonisation levels and acts systemically to suppress colonisation rates (Balzergue et al. 2013; Breuillin et al. 2010). Recent studies in rice, *Lotus japonicus* and *M.truncatula* indicate phosphorous suppresses mycorrhizae through an interaction of SPX phosphate sensors with phosphate starvation response (PHR) transcription factors (Das et al. 2022; Shi et al. 2021; Wang et al. 2021b). CLE peptides have also been proposed as potential regulators of phosphate control of mycorrhizal colonisation, as high phosphate has been shown to induce the expression of specific CLE peptides, some of which are also induced by mycorrhizal colonisation (Funayama-Noguchi et al. 2011; Handa et al. 2015; Karlo et al. 2020; Müller et al. 2019).However, the fact that both legume and non-legume mutants disrupted in CLE signalling pathway elements all still display strong mycorrhizal suppression in response to phosphate (Foo et al. 2013; Müller et al. 2019; Wang et al. 2021a; Wyss et al. 1990) indicates this pathway does not appear to play a particularly important role. A more direct approach to examining CLE function in phosphate response is therefore required.

In this study we examined a range of CLE peptides in the non-legume model tomato and identify two CLE peptides, *Sl*CLE10 and *Sl*CLE11, with important roles in limiting mycorrhizal colonisation of the root. Mutant and transgenic studies indicated a more important role for *Sl*CLE11. We also examined the role of the CLE receptors *Sl*FAB and *Sl*CLV2 in *Sl*CLE11 perception and the role of *Sl*CLE11 in phosphate suppression of mycorrhizal colonisation.

## Results

### Characterisation of CLE peptides in tomato

Recent analysis has identified 52 putative *SlCLE* in tomato, with at least 43 expressed (Carbonnel et al. 2022). Phylogenetic analysis of all 52 *Sl*CLE peptides and *Mt*CLE peptides (Hastwell et al. 2017) was undertaken (Fig 1). Interestingly, all the CLE peptides from *M.truncatula* with a characterized function in the autoregulation of nodulation (AON) or autoregulation of mycorrhization (AOM) processes belong to the same clade, and many of them are reported to be transcriptionally regulated not only by symbiosis (nodulation and AM) but also by nitrogen and phosphorous (Fig 1). This clade also contain a number of *Sl*CLE peptides.

**Figure 1.**
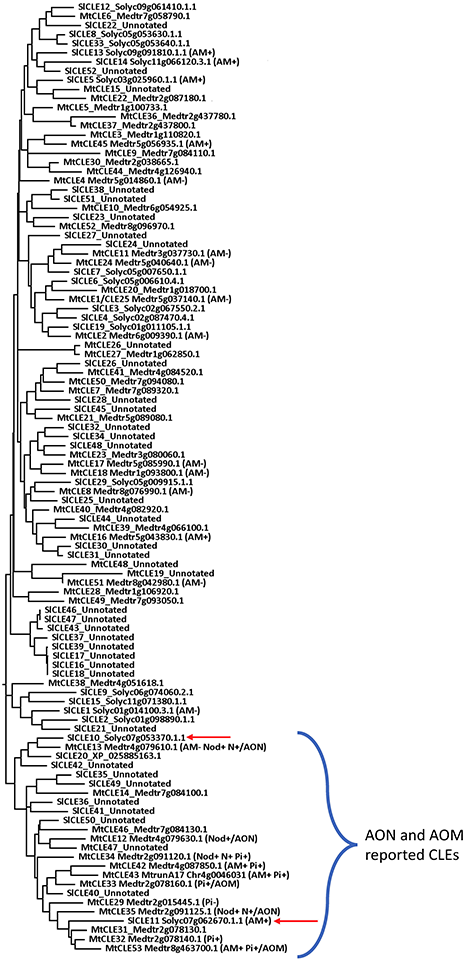
Phylogenetic tree of CLE peptide families from tomato and *Medicago truncatula*. Amino acid sequences from the CLE peptide families in tomato (Carbonnel et al. 2022) and *Medicago truncatula* (Hastwell et al. 2017), were aligned using the MUSCLE algorithm of MEGA11 software (Tamura et al. 2021). Phylogenetic relationships were determined with MEGA11 to create a maximum-likelihood (ML) tree using Jones–Taylor–Thornton (JTT) as the amino acid substitution model and the nearest-neighbor-interchange (NNI) heurist method to improve the likelihood of the tree. The partial deletion (95%) mode was used for the treatment of gaps and missing data. 100 bootstrap replications were performed. CLE peptides whose gene expression is induced (+) or repressed (−) during mycorrhization (AM), nodulation (Nod), high phosphate treatment (P) or nitrate treatments (Karlo et al. 2020; Lebedeva et al. 2020; Mens et al. 2020; Mortier et al. 2010; Müller et al. 2019) are indicated between brackets. Those CLE peptides with a reported function in the autoregulation of nodulation (AON) or autoregulation of mycorrhization (AOM) (Imin et al. 2018; Lebedeva et al. 2020; Mens et al. 2020; Mortier et al. 2010; Müller et al. 2019) are also indicated, and all of them belong to the same clade. *Sl*CLE10 and *Sl*CLE11 peptides, that are functionally characterized in this study, are labelled with a red arrow. The tree is shown with bootstrap confidence values superior to 1.

At the time of the study, fifteen genes coding for CLE peptides had been identified in tomato (SlCLE1-SlCLE15; Zhang et al. 2014) and these were the focus of gene expression studies (Table 1). We examined the effect of nitrate and phosphate levels on the expression of *SlCLE* genes in the absence of mycorrhizal colonisation (Table 1). Only *SlCLE2, SlCLE10* and *SlCLE14* were found to be significantly up-regulated by high nitrate, while none were significantly up-regulated by phosphate, although *SlCLE10* and *SlCLE11* were somewhat upregulated by high phosphate (Table 1). The influence of mycorrhizal colonisation on the expression of CLE family genes was investigated using the RNA-seq data from the NCBI database (Accession No.PRJNA509606 and PRJNA523214; Ho-Plagaro et al. 2020). Four *SlCLE* genes (*SlCLE 5,11,13* and *14*) were found to be significantly induced by mycorrhizal colonisation, with *SlCLE11* displaying a nearly five-fold induction in expression compared to uncolonised plants (Table 1). Further, in *SlDLK2* silenced mycorrhizal roots, which are characterized by an increased number of highly branched and transcriptionally active arbuscules (Ho-Plagaro et al. 2020), *SlCLE11* gene expression was increased approximately 95-fold compared to uncolonised empty vector control plants (Table 1). Two *SlCLE* genes were significantly suppressed in response to mycorrhizal colonisation (*SlCLE1* and *SlCLE6*).

**Table 1.** Relative expression of *SlCLE* genes in non-mycorrhizal (NM) plants in response to nitrate (LN, low nitrate; MN, medium nitrate; HN, high nitrate) and phosphate (LP, low phosphate; MP, medium phosphate; HP, high phosphate) and in response to arbuscular mycorrhizal (AM) colonisation with *R. irregularis* in control and *SlDLK2* RNAi plants. *n.d., not detected*, gene expression was below detection limit. *n.t., not tested*, the expression of this gene not tested. Within a gene and experiment, values that were significantly different indicated with different letters (ANOVA, followed by Tukeys HSD test, P<0.05) or t-test (P<0.05 *, P< 0.01 **).

| Gene expression | Nitrate |  | Phosphate |  | AM (compared to NM control) | AM SIDLK2 RNAi (compared to NM control) |
| --- | --- | --- | --- | --- | --- | --- |
| *SICLE1*<br>*Solyc01g014100* | LN | *nd* | LP | *nd* | 0.16* | 0.01** |
|  | MN | *nd* | MP | *nd* |  |  |
|  | HN | *nd* | HP | *nd* |  |  |
| *SICLE2*<br>*Solyc01g098890* | LN | 0.79a | LP | 0.75 | 1.24 | 1.08 |
|  | MN | 1.00a | MP | 1.00 |  |  |
|  | HN | 1.54b | HP | 1.07 |  |  |
| *SICLE3*<br>*Solyc02g067550* | LN | 0.84 | LP | *nt* | 0.54 | 0.61 |
|  | MN | 1.00 | MP | *nt* |  |  |
|  | HN | 1.09 | HP | *nt* |  |  |
| *SICLE4*<br>*Solyc02g087470* | LN | 0.96 | LP | 1.22 | 1.22 | 1.04 |
|  | MN | 1.00 | MP | 1.00 |  |  |
|  | HN | 1.22 | HP | 1.73 |  |  |
| *SICLE5*<br>*Solyc03g025960* | LN | 1.01 | LP | *nt* | 2.04* | 2.26* |
|  | MN | 1.00 | MP | *nt* |  |  |
|  | HN | 1.26 | HP | *nt* |  |  |
| *SICLE6*<br>*Solyc05g006610* | LN | 1.02 | LP | 3.92 | 0.65 | 0.26* |
|  | MN | 1.00 | MP | 1.00 |  |  |
|  | HN | 1.29 | HP | 6.92 |  |  |
| *SICLE7*<br>*Solyc05g007650* | LN | 0.76 | LP | 1.24 | 1.29 | 1.22 |
|  | MN | 1.00 | MP | 1.00 |  |  |
|  | HN | 0.97 | HP | 1.92 |  |  |
| *SICLE8*<br>*Solyc05g053630* | LN | 0.79 | LP | *nt* | 0.00 | 0.00 |
|  | MN | 1.00 | MP | *nt* |  |  |
|  | HN | 0.96 | HP | *nt* |  |  |
| *SICLE9*<br>*Solyc06g074060* | LN | *nt* | LP | *nt* | 0.99 | 0.90 |
|  | MN | *nt* | MP | *nt* |  |  |
|  | HN | *nt* | HP | *nt* |  |  |
| *SICLE10*<br>*Solyc07g053370* | LN | 0.61a | LP | 1.70 | 1.54 | 1.14 |
|  | MN | 1.00a | MP | 1.00 |  |  |
|  | HN | 9.45b | HP | 4.97 |  |  |
| *SICLE11*<br>*Solyc07g062670* | LN | 0.90 | LP | 1.54 | 4.93* | 94.91** |
|  | MN | 1.00 | MP | 1.00 |  |  |
|  | HN | 0.76 | HP | 4.68 |  |  |
| *SICLE12*<br>*Solyc09g061410* | LN | 1.30 | LP | 1.21 | 1.18 | 2.15 |
|  | MN | 1.00 | MP | 1.00 |  |  |
|  | HN | 1.11 | HP | 1.42 |  |  |
| *SICLE13*<br>*Solyc09g091810* | LN | *nd* | LP | *nt* | 3.00* | 2.92 |
|  | MN | *nd* | MP | 1.00 |  |  |
|  | HN | *nd* | HP | 1.30 |  |  |
| *SICLE14*<br>*Solyc11g066120* | LN | 0.82a | LP | *nt* | 2.02* | 1.50 |
|  | MN | 1.00a | MP | *nt* |  |  |
|  | HN | 1.61b | HP | *nt* |  |  |
| *SICLE15*<br>*Solyc11g071380* | LN | *nt* | LP | *nt* | 0.00 | 1.14 |
|  | MN | *nt* | MP | *nt* |  |  |
|  | HN | *nt* | HP | *nt* |  |  |

*Sl*CLE10 and *Sl*CLE11 were selected for further study. This was due to *Sl*CLE10 and *Sl*CLE11 homology with *Mt*CLE peptides with roles in symbioses and/or nutrient response (Fig 1) and the fact that *SlCLE11* expression was induced by AM colonisation and *SlCLE10* expression was induced by nitrate (Table 1). The expression of the *SlCLE10* and *SlCLE11* genes was analysed in different organs of non-inoculated tomato plants by RT-qPCR (Suppl Fig 1A, B). *SlCLE10* was most highly expressed in roots, while *SlCLE11* was highly expressed in fruits, reaching the peak of expression in fruits turning red (Suppl Fig 1B). Similar results were obtained by Zhang et al. (2014) from tomato cv. ‘Heinz-1706’, although they observed a peak in *SlCLE11* expression at the earliest (initial green fruits) and the latest (red fruits) stages of fruit development, possibly due to differences in the growth conditions or the tomato variety. To examine a possible link between *SlCLE* expression and ethylene-induced processes such as fruit ripening, we analysed the expression of *SlCLE10* and *SlCLE11* in tomato roots in response to ethephon application. *SlCLE11* expression displayed a significant but transient increase in expression 12h after ethephon treatment compared to control plants (Suppl Fig 1C) and this was validated in an independent experiment (data not shown). No change in *SlCLE10* expression was observed in response to ethephon (data not shown).

### Sl*CLE10 and* Sl*CLE11 suppress mycorrhizal colonisation*

To examine if *Sl*CLE10 and/or *Sl*CLE11 regulate mycorrhizal colonisation, lines with CRISPR-induced mutations were generated for each peptide (Suppl Fig 2). The *cle10* mutant line has a large deletion resulting in a predicted *Sl*CLE10 protein missing 30 amino acids from the middle of the polypeptide and introduces a premature stop codon before the CLE motif. The *cle11* mutant has a 3 bp deletion leading to deletion of an isoleucine (32) and also 1bp deletion leading to a premature stop codon before the CLE motif. Mycorrhizal colonisation was not significantly different in *cle10* mutant plants compared to wild type lines in two independent experiments (Fig 2 A,C). In contrast, a significant increase in all fungal structures and also in arbuscules was seen in the roots of *cle11* mutant plants compared to wild type lines (approximately 30% increase; Fig 2B). A significant increase in the amount of the root colonised and the amount of the root containing arbuscules was also observed in *cle11* mutants compared to wild type an independent experiment and this increase in arbuscules was similar to that observed in *fab* mutants (Fig 2C). Plants disrupted in both peptides, *cle10 cle11* double mutant, displayed a similar increase in arbuscule colonisation as observed in *cle11* mutants, although there was only a small but no significant increase in total colonisation of *cle10 cle11* double mutant roots compared to wild type (Fig 2C).

**Figure 2.**
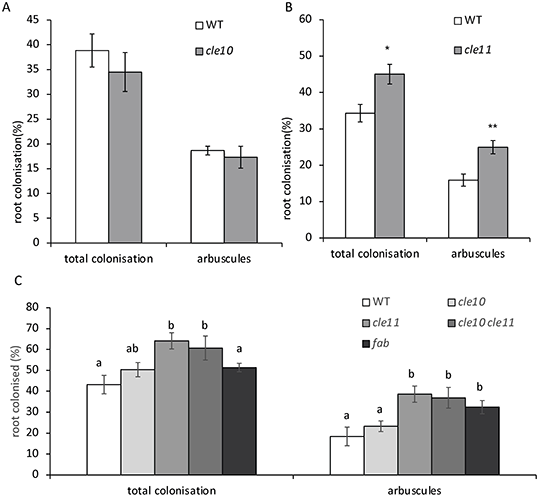
Mycorrhizal colonisation in wild type (WT) plants and (A) *cle10* and (B) *cle11* and (C) *cle10 cle11* double mutant plants. Data shown are mean ± SE, (n=7-9). For (A, B) values that are significantly different to WT, * P<0.05, ** P<0.01, for (C) within a parameter different letters indicate values that are significantly different as assessed by Tukey’s HSD test (P < 0.05).

Another approach to examine the influence of *Sl*CLE10 and *Sl*CLE11 on mycorrhizal colonisation is to overexpress each peptide in hairy-root transformed tomato. Overexpression of *SlCLE11* in wild type pea roots led to a significant increase in *SlCLE11* expression (Fig 3A) and a significant decrease in mycorrhizal colonisation in transformed roots compared to roots transformed with empty vector control (GFP+; transformed roots detected by expression of GFP; Fig 3B). Total fungal colonisation was reduced approximately 3-fold, while amount of root colonised by arbuscules was reduced approx. 4-fold. Interestingly, overexpression of *SlCLE11* did not influence mycorrhizal colonisation in non-transformed (-GFP) roots of the same plant, suggesting the effect of *Sl*CLE11 was not systemic (Fig 3B). It is also important to note that transformation *per se* did not influence colonisation rates, as there was no difference in colonisation rates between transformed (+GFP) and untransformed roots (-GFP) on the same empty vector (EV) control plants (Fig 3B). In an independent experiment, a similar significant decrease in total colonisation and arbuscules was observed in the roots of *SlCLE11* overexpressing roots compared to plants expressing empty vector control colonised with *R.irregularis* or *F.mosseae* (Suppl Fig 3A, B). The influence of overexpression of *SlCLE10* was also examined in wild type pea roots (Fig 4). Elevated expression of *SlCLE10* (Fig 4A) led to a significant approx. 2-fold reduction in total fungal colonisation and amount of root colonised with arbuscules, compared to plants transformed with empty vector control (Fig 4B).

**Figure 3.**
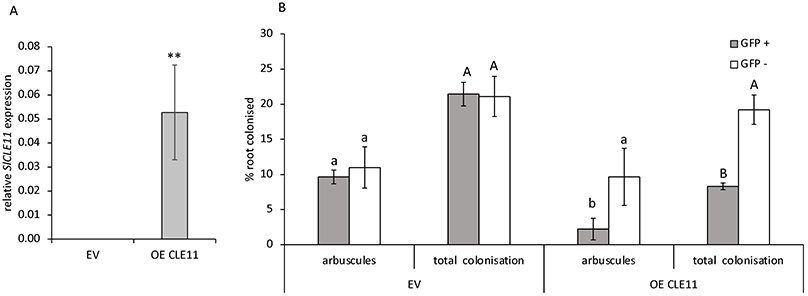
*SlCLE11* overexpression (OE) and empty vector (EV) transformed (GFP+) and non-transformed (GFP-) wild type tomato roots. A. Relative *SlCLE11* gene expression in transformed roots. B. Mycorrhizal colonisation. Data shown are mean ± SE, (n = 5-8). A. Values that are significantly different to WT,** P < 0.01. **B**. Within a parameter, different letters indicate values that are significantly different as assessed by Tukey’s HSD test (P < 0.05).

**Figure 4.**
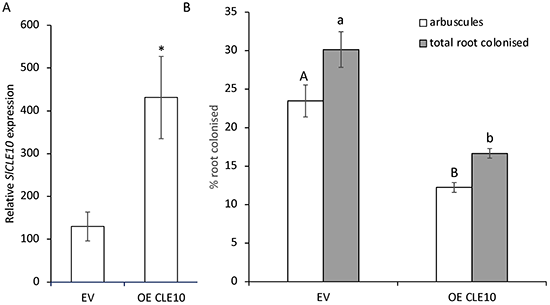
*SlCLE10* overexpression (OE) and empty vector (EV) transformed wild type tomato roots. (A) *SlCLE10* gene expression (B) Mycorrhizal colonisation. For (A) values that are significantly different to WT,** P < 0.01, for (B)Within a parameter different letters indicate values that are significantly different as assessed by Tukey’s HSD test (P < 0.05).

### Sl*CLE11 does not act via LRR receptors* FAB *and* CLV2 *and is not required for phosphate regulation of mycorrhizal colonisation*

Given the strong phenotype of *cle11* mutants and roots overexpressing *SlCLE11*, further studies were undertaken to examine the function of the *SlCLE11* gene. Two putative receptors, FAB and *Sl*CLV2, that negatively regulate mycorrhizal colonisation have been identified in tomato (Wang, CL *et al*., 2018; Wang, C *et al*., 2021). To test if these may be the receptor(s) for the *Sl*CLE11 peptide, the *SlCLE11* gene was overexpressed in *fab*, *clv2-5* mutant and wild type tomato roots (Fig 5). *SlCLE11* expression was similar in all genotypes transformed with the empty vector and was elevated approx. 20-fold in all genotypes in *35S∷SlCLE11* transformed roots (Fig 5A). As previously observed in wild type plants (Fig 3; Suppl Fig 3), overexpression of *SlCLE11* significantly reduced total fungal colonisation and amount of root colonised with arbuscules by approximately 3- to 4-fold compared to the EV wild type controls (Fig 5B). A similar approx. 3- to 4-fold reduction in the amount of root colonised with any fungal structure or with arbuscules alone was observed when *SlCLE11* was overexpressed in *fab* and *clv2-5* mutant roots compared to EV controls of each genotype, indicating that *FAB* and *SlCLV2* are not required for the *SlCLE11* response. It is interesting to note that in this and several other transgenic experiments (data not shown), AM colonisation was not significantly elevated in *fab* or *clv2* mutant plants compared to wild type, in contrast to significant elevation in AM colonisation in these genotypes when not transformed (e.g. Fig 2, 6, 7; Wang et al. 2021a; Wang et al. 2018). This may be due to altered root growth dynamics in transgenic tomato cuttings influencing the rate of root colonisation and possibly the influence of *FAB* and/or *SlCLV2* on colonisation in control plants.

**Figure 5.**
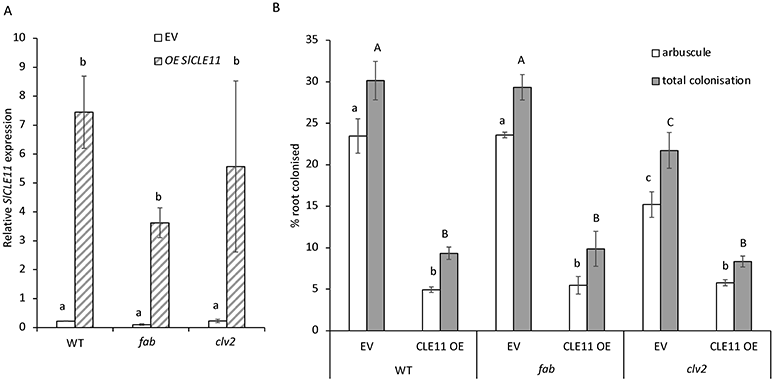
*SlCLE11* overexpression (OE) and empty vector (EV) transformed wild type (WT), *fab* and *clv2-5* mutant tomato roots. (A) Relative *SlCLE11* gene expression in transformed roots. (B) Mycorrhizal colonisation. Data shown are mean ± SE, (n = 5-10). Within a parameter, different letters indicate values that are significantly different as assessed by Tukey’s HSD test (P < 0.05).

**Figure 6.**
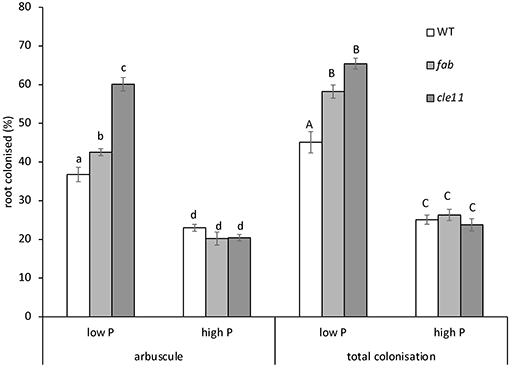
Phosphate control of mycorrhizal colonisation in WT, *fab* and *Slcle11* plants. Plants received modified LANS with 0.05 (low P) or 5mM (high P) NaH_2_PO_4_. Values are mean + standard error (n=6). Within a parameter, different letters indicate values that are significantly different as assessed by Tukey’s HSD test (P < 0.05).

**Figure 7.**
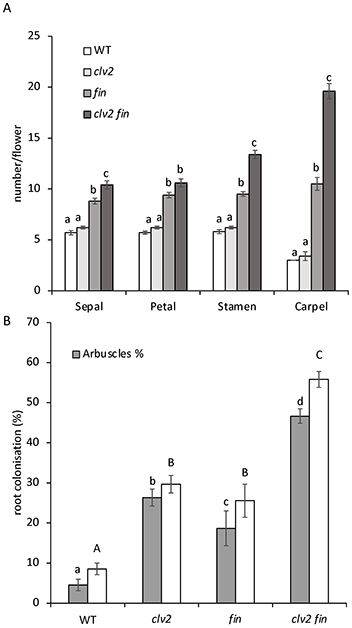
Development of wild type (WT), *fin, clv2* and *fin clv2* double mutant plants. A. Floral organ number, B. Mycorrhizal colonisation. Values are mean + SE, for A (n = 3), for B (n = 8-10). Within a parameter, different letters indicate values that are significantly different as assessed by Tukey’s HSD test (P < 0.05).

We also examined if phosphate regulation of mycorrhizal colonisation requires *Sl*CLE11. We monitored AM colonisation in wild type, *fab* mutants and *cle11* mutants grown under low and high phosphate (Fig 6). As previously reported, *fab* and *cle11* mutants displayed elevated colonisation rates compared to wild type plants (Fig 2; Wang et al. 2021a). High phosphate levels inhibit AM colonisation of roots in wild type and *fab* mutant tomato plants compared to plants grown with low phosphate (Fig 6). Similarly, AM colonisation was suppressed in *cle11* mutant plants grown under high phosphate compared to low phosphate controls. Indeed *cle11* mutants grown under high phosphate had similar levels of colonisation to wild type plants grown under high phosphate (Fig 6).

Previous double mutant studies have indicated that FAB and *Sl*FIN have additive effects on shoot meristem, flower and fruit development in tomato (Xu et al. 2015). We generated a *fin clv2* double mutant and examined the effect on shoot development and mycorrhizal colonisation compared to the single mutant parents and wild type plants (Fig 7). In this study, *clv2* mutants did not display any significant changes to floral organ number or shoot morphology, while as observed previously, *fin* mutants displayed significantly more floral organs and an increase in number of flowers in each inflorescence compared to wild type plants (Fig 7 A, Suppl Fig 4; Xu *et al*., 2015). The double mutant *clv2 fin* plants displayed a clear additive phenotype, with a significant increase in the number of sepals, stamens and carpels compared to *fin* single mutant plants (Fig 7A) and an increase in inflorescence complexity and stem fasciation (Suppl Fig 4). Both *clv2* and *fin* mutants exhibited a significant increase (3- to 4-fold) in AM colonisation, including all fungal structures and arbuscules, compared to wild type plants (Fig 7B). The *clv2 fin* double mutant plants displayed a clear additive phenotype, with approx. 7-fold increase in total root colonised and approx. 10-fold increase in the amount of root colonised by arbuscules compared to wild type, significantly higher than either single mutant parent (Fig 7B).

## Discussion

In this study we identified two tomato CLE proteins with roles in the suppression of arbuscular mycorrhizal colonisation, *Sl*CLE10 and *Sl*CLE11. *Sl*CLE11 appears to play a key role in regulating colonisation, as disruption of *Sl*CLE11 led to a significant increase in colonisation and overexpression significantly suppressed mycorrhizal colonisation (Fig 2, 3, Suppl Fig 3). In contrast, although overexpression of *Sl*CLE10 suppressed colonisation, mutation of *Sl*CLE10 did not influence colonisation rates (Fig 2, 4). Furthermore, double mutant *cle10 cle11* plants displayed a similar increase in colonisation to *cle11* single mutant plants (Fig 2), indicating loss of function of *cle10* had little influence on colonisation. Taken together, these results suggest that although ectopic expression of *SlCLE10* influenced colonisation, this gene is likely to play only a minor or redundant role in regulating colonisation rates. In contrast, *SlCLE11* appears to play a prominent role in regulating colonisation rates and this is supported by the fact that expression of *SlCLE11* is dramatically induced in roots colonised with mycorrhizal fungi (Table 1). Transgenic studies with composite plants indicate that the *SlCLE11* peptide acts locally to suppress colonisation, with no evidence that the peptide acts systemically (Fig 3). It is interesting to note that ethylene, which can suppress mycorrhizal colonisation in tomato and other species (e.g. Torres de Los Santos *et al*., 2014; Foo *et al*., 2016), also induces the expression of *SlCLE11* (Suppl Fig 2), suggesting a potential interaction between ethylene and *SlCLE11* during mycorrhizal interactions.

*SlCLE11* is the most closely related of all of the *SlCLE* genes to *MtCLE53* (Fig. 1), which encodes a CLE peptide with a major role in regulating colonisation *in M.truncatula* via the *Mt*SUNN CLV1 receptor (Karlo et al. 2020; Müller et al. 2019). The CLV1 orthologue in tomato, FAB also plays an important role in suppressing mycorrhizal colonisation in tomato (Fig 5, 6; Wang et al. 2021a) and has been demonstrated to perceive CLE peptides *in vivo* (Xu et al. 2015). However, we found no evidence that FAB is required to perceive *Sl*CLE11, as ectopic expression of *SlCLE11* still suppressed mycorrhizal colonisation in the *fab* mutant (Fig 5). Similarly, although *SlCLV2* is another CLE receptor known to suppress mycorrhizal colonisation (Fig 7; Carbonnel et al. 2022; Wang et al. 2021a), *Sl*CLE11 does not appear to interact with this receptor to suppress colonisation (Fig 5). Previous studies have found additive shoot phenotypes in the *fab fin* double mutant, suggesting these proteins act in somewhat parallel pathways (Xu et al. 2015). Similarly, we found that *clv2 fin* double mutants also display additive shoot and mycorrhizal colonisation phenotypes compared to the single mutant *clv2* and *fin* parents (Fig 7, Suppl Fig 4). Future studies could explore the perception pathway of *Sl*CLE11 and seek to identify the *Sl*CLE peptides that act through FAB, *Sl*CLV2 and FIN proteins to suppress colonisation. This might include other *Sl*CLE peptides related to symbioses CLEs (Fig 1) and/or other *Sl*CLEs whose expression is induced by mycorrhizal colonisation (Table 1).

Although *Sl*CLE11 plays an important role in suppressing mycorrhizal colonisation, studies with *cle11* mutant did not indicate any involvement in suppressing colonisation in response to phosphate (Fig 6). This suggests that as observed in legumes (outlined in the introduction), there is little evidence that CLE peptide signalling plays a role in phosphate control of mycorrhizal colonisation. In contrast, some of the CLE signalling elements are required for nitrogen suppression of mycorrhizal colonisation levels in tomato (Wang et al. 2021a) and future studies could explore the *Sl*CLE peptides involved. *Sl*CLE10 is a possible candidate, as expression of the *SlCLE10* gene was significantly elevated by high nitrogen (Table 1). It is important to note that all experiments presented in this study were grown under relatively high nitrogen but low phosphate levels. Given that in tomato phosphate is a much more potent suppressor of mycorrhizal colonisation than nitrate, this may have masked any difference between wild type and *cle10* mutants. In addition, the phenotype of single *cle* mutants could be masked by compensation, given that both active and passive compensation of CLE paralogues operate in the control of the shoot apex size in tomato, Arabidopsis and maize (Nimchuk et al. 2015; Rodriguez-Leal et al. 2019). Future studies could explore if similar compensation operates in CLE regulation of symbioses.

In conclusion, our results show that the control of arbuscular mycorrhizal colonisation by CLE peptide signalling is conserved across legume and non-legume species. This study has identified the first non-legume CLE peptides involved in this pathway and has revealed that receptors in addition to CLV1 and CLV2 orthologues must play a role in this important control system.

## Materials and Methods

### Plant material, growth conditions and mycorrhizal studies

Several studies were conducted with the tomato *Solanum lycopersicum* wild type cv. Money Maker (Table 1 mycorrhizal studies; Suppl Fig 1 and 3). All other experiments employed the tomato wild type cv. M82, and mutants on this background; *fab*, *Slclv2-5, fin-n2326* (Wang et al. 2021a; Xu et al. 2015), *Slcle10* and *Slcle11* (described in this manuscript). Mutations introduced in *cle10* and *cle11* are outlined in Suppl Fig 2. Double mutants disrupted in both *cle10* and *cle11* and also *Slclv2* and *fin* were each generated by crossing the parental mutants and selecting the double mutants in the F2 using PCR or high resolution melt PCR (primers outlined in Suppl Table 1).

Unless otherwise stated mycorrhizal inoculum was live corn pot culture originally inoculated with spores of *Rhizophagus irregularis* (INOQ Advantage, INOQ GMBH, Germany), grown under glasshouse conditions that received modified Long Ashton nutrient solution (Hewitt 1966) containing 3.7mM KNO_3_ and 0.5mM NaH_2_PO_4_ twice a week (1 - 3 weeks after transplanting) and three times a week (from 3 weeks after transplanting). For phosphate response experiment (Fig 6), plants received modified LANS with 0.05 (low P) or 5mM (high P) NaH_2_PO_4_. The inoculum contained colonised root segments, external hyphae and spores and for experiments the growth substrate (80%) was mixed with whole inoculum (20%). Tomato seeds were germinated in potting mix and transplanted two weeks after sowing into 2L pots containing a 1:1 mixture of vermiculite and gravel (plus inoculum), topped with vermiculite. Plants were grown in controlled glasshouse (25 °C day/20 °C night, 18 hour photoperiod). For mycorrhizal studies outlined in Table 1 and studies in Suppl Fig 3 tomato plants were inoculated with *R. irregularis* (DAOM 197198) or *F.mosseae* (Schußler and Walker) and were grown in a growth chamber (24 °C day/20°C night,16 hour photoperiod) in 0.5L pots containing an autoclave-sterilized mixture of expanded clay, washed vermiculite, and coconut fiber (2:2:1, by volume). For *R. irregularis* inoculum, a piece of monoxenic culture in Gel-Gro medium produced according to Chabot et al. (1992), containing 50 spores of *R. irregularis* and infected carrot roots was used. For the *F. mosseae*-inoculated treatment, inoculum was obtained from the EEZ germplasm collection and the fungus was propagated using sorghum (*Sorghum bicolor*) as the host. The infected roots, hyphae, spores and substrates were collected and used as inoculum (40 g per pot).

Roots were stained with trypan blue or the ink and vinegar method (Vierheilig et al. 1998). Mycorrhizal colonisation of roots was scored according to McGonigle et al. (1990), where 150 intersects were observed from 25 root segments per plant. The presence of arbuscules, vesicles and intraradical hyphae at each intersects was scored separately. The total colonization of mycorrhizae was calculated as the percentage of intersects that have presence of any fungal structures and arbuscule frequency was calculated from the percentage of intersects that contained arbuscules.

### Gene expression studies

For the DLK2 RNAi study, the results from previously published RNA-seq data from the NCBI accessions PRJNA509606 and PRJNA523214 are presented (Ho-Plagaro et al. 2020). For gene expression in different tissues, non-inoculated wild type tomato plants were grown in a glasshouse (light:dark regime of 16:8h at 24:18°C) in 10 L pots containing peat and watered with tap water as needed. Leaves, stems, roots, as well as flowers and fruits at different developmental stages were collected. For the ethephon experiment wild type tomato plants were grown in 200 ml pots filled with a mixture of silicate sand, peat, soil, and vermiculite (1:1:1:1 v:v:v:v). Forty days later, 25 mL of 75 μM ethephon was applied to roots and the root tissue was harvested 12, 24 and 48 h later for gene expression analysis. For gene expression studies under various doses of N or P, tomato wildtype plants were grown with no mycorrhizal inoculum with various nutrient treatments and harvested 6-7 weeks after sowing. For the N experiment, plants received modified Long Ashton Nutrient Solution containing 5mM NaH_2_PO_4_ and 0.625, 2.5 or 10mM KNO_3_. For the P experiment plants received modified LANS with 3.7mM KNO_3_ and 0.05, 0.5 or 5mM NaH_2_PO_4_. For all gene expression studies roots were cut into 1 - 2cm segments and the roots from three different plants were mixed well to form one biological replicate. The total RNA was extracted from ground root samples using an ISOLATE II Plant Mini Kit (Bioline, Australia), cDNA was synthesised from 1ug of RNA using a SensiFAST cDNA Synthesis Kit (Bioline, Australia) and real time PCR reactions were completed in duplicate for each sample using a SensiMix SYBR (Bioline); 95◻ for 10 min, 50 cycles of 95◻ for 5s, 60 or 59◻ for 40s. The *SlEF1 (Elongation factor 1*α) and *SlCAC* (Clathrin adaptor complexes) genes were used as control (Lacerda *et al*., 2015). Standard curves were created for each gene using serially diluted plasmids containing cloned fragments of each amplicon. The average concentration of technical replicates was calculated. The relative gene expression of biological replicates was determined by comparing the concentration of the gene of interest with the concentration of the two control genes for this sample.

### CRISPR-Cas9 gene editing and transformation

CRISPR–Cas9 mutagenesis of tomato was performed as described previously (Brooks et al. 2014; Kwon et al. 2022; Van Eck et al. 2019). Briefly, guide RNAs (gRNAs) were designed using the CRISPRdirect software (Naito et al. 2015) (https://crispr.dbcls.jp/) and a binary vector (pAGM4723) was built with a Cas9 driven by cauliflower mosaic virus 35S promoter and gRNAs driven by the *Arabidopsis* U6 promoter through Golden Gate cloning (Weber et al. 2011; Werner et al. 2012). The final binary plasmids were introduced into the tomato cultivar M82 by *Agrobacterium tumefaciens*–mediated transformation as described previously (Van Eck et al. 2019). First-generation (T_0_) transgenic plants were transplanted in soil and grown in a greenhouse under long-day conditions (16 h light, 26–28 °C/8 h dark, 18–20 °C; 40–60% relative humidity). All T_0_ transgenic plants were backcrossed to the M82 and genotyped by PCR (35S-seq-F CTGACGTAAGGGATGACGCAC, Cas9-CATCTCATTACTAAAGATCTCC) and sprayed with 400 mg/L kanamycin to confirm absence of the transgene. For genotyping of CRISPR-generated mutations, genomic DNA was extracted by standard CTAB protocol and each targeted gene was amplified by PCR and sequenced via Sanger sequencing. Stable non-transgenic, homozygous plants were used for all experiments.

### Hairy root transformation

To generate the binary vectors, specific promoters and genes were inserted in PK7WG2D vector using gateway technology. Briefly, PCR fragments for genes were generated using primers containing the attB recombination site and inserted into the entry vector and subsequently, the destination vector using two recombination reactions, following the instructions of the Gateway cloning system (Invitrogen Life Technologies). The *SlCLE10* and *SlCLE11* coding regions were amplified from tomato cDNA obtained from M82 roots and confirmed by sequencing and expression was driven by 35S promoter. Plasmids containing the ccB gene (empty vector; EV) were used as a negative control. Unless otherwise stated, *Agrobacterium rhizogenes* (K599) was used to transform tomato plant roots on non-transformed shoots, following a method previously established for pea (Clemow et al. 2011) with some modifications. *A.rhizogenes* was grown in luria broth with appropriate antibiotics, for 24 hours at 28°C at 170rpm. Shoot cuttings were taken from tomato plants approx. 21 days after sowing and placed in a rock wool previously inoculated with 40mL of bacterial suspension (~OD 0.4). Plants were kept in mini glasshouses for 24 hours with lid holes closed, then for an additional 24 hours without lids to promote wilting of the shoots, watered and covered with lids again for 24 hours, in low light. Plants were kept for an additional 10 days at 21°C and 18 hours light. Rockwool was removed and plants were transferred to 1L pots containing inoculum as outlined above. For studies presented in Suppl Fig 3, hairy root transformation was performed according to Ho-Plagaro et al. (2018) using *A.rhizogenes* MSU440 harbouring the pUBIcGFP-DR (Kryvoruchko et al. 2016). Plants were harvested 4-5 weeks after transplanting. Roots of 10 to 20 plants per treatment were washed and each individual root generated from the callus was dissected and checked for GFP fluorescent protein as a reporter for transformation. Transformed and non-transformed roots were harvested separately, with roots cut into 1.5cm segments, a small portion collected for RNA extraction (stored at −80°C), with the remaining roots were stored in 50% EtOH for later staining (outlined above).

## Data availability

Plant material and constructs are available upon request.

## Acknowledgements

We thank Zachary Lippman (Cold Spring Harbour Laboratory) for supplying the tomato mutants and Tracey Winterbottom and Sarah Kane for assistance with plant husbandary.

## Author contributions

E.F. conceived the project, with input from J.B.R., T. H-P. and J.M.G.G. C-T.K. generated the CRISPR mutant lines. K.V. optimised the tomato transformation system for studies undertaken at UTAS. K.W., C.W., T.H-P., K.V., M.I.T-N., A.C-L., J.S. and E.F. performed the experiments and analysis. E.F. wrote the manuscript with input from all authors.

