## Supplementary material for "CLE11 and CLE10 Suppress Mycorrhizal Colonisation in Tomato": Suppl Figs

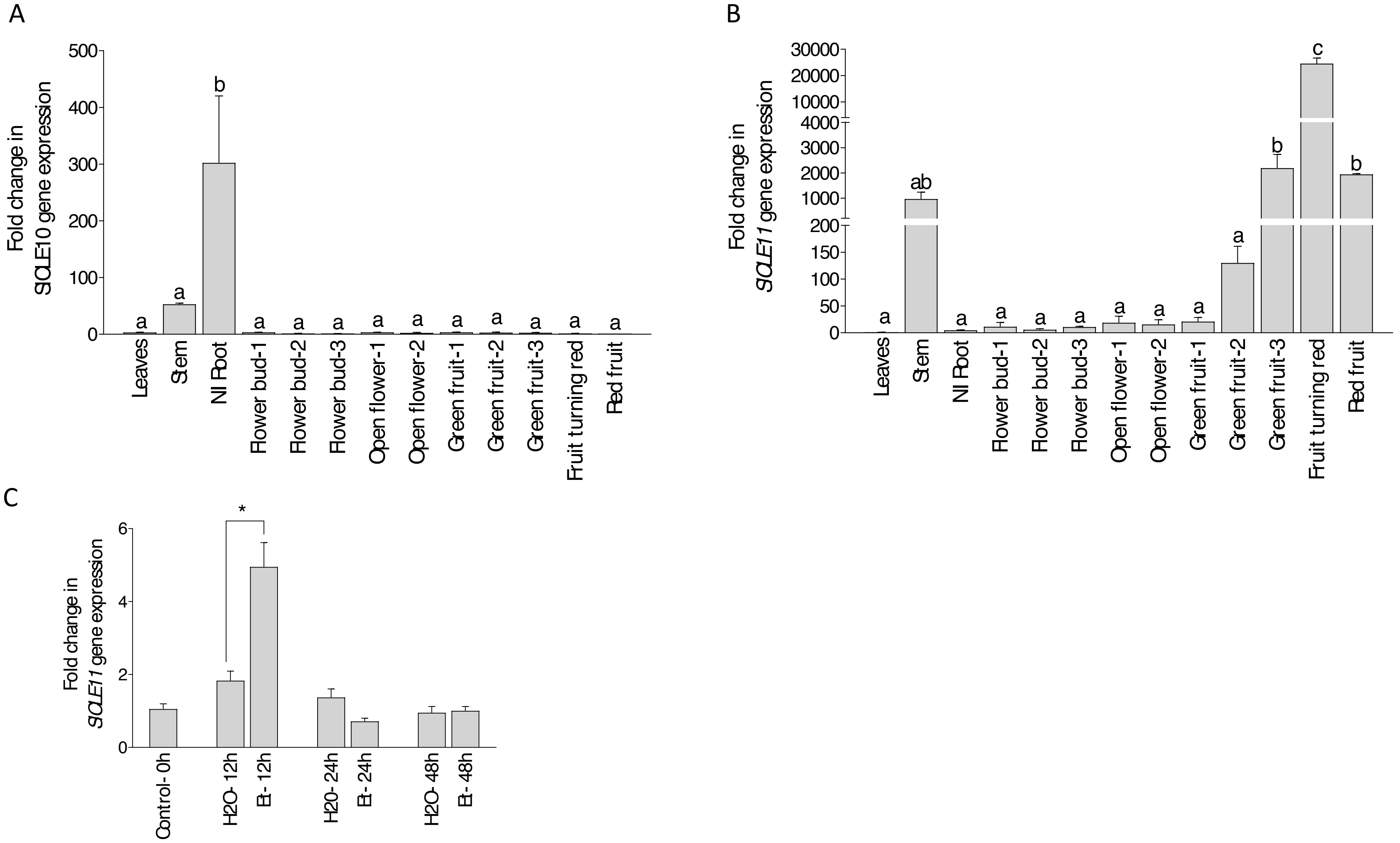


**Suppl Figure 1. *SlCLE10* and *SlCLE11* gene expression in different tomato organs.** In tomato wild type, *SlCLE10* (A) and *SlCLE11* (B) gene expressions were measured by RT-qPCR in leaves, stems, non-inoculated (NI) roots; initial, young and mature flower buds (flower buds-1, -2 and -3, respectively); young and mature open flowers (open flowers -1 and -2, respectively); green fruits at increasing developmental stages (green fruits-1, -2, -3), fruits turning red, and mature fruits in red. Data represent the mean ± SE (n=3). RT-qPCR data represents the relative expression with respect to the organ showing the lowest expression level (red fruits in "A" and leaves in "B"), in which gene expression was designated as 1. Significant differences ($P < 0.05$) observed by the LSD multiple comparison test are labelled with different letters. © tomato plantlets treated with 70 µM ethephon ("Et") for 12, 24 and 48 hours. Values correspond to mean ± SE (n=3). RT-qPCR data represents the relative expression with respect to the control plants prior to treatment (0h), in which gene expression was designated as 1. Significant differences (Student's t-test) between the ethephon-treated plants ("Et") and the $H_2O$-treated plants ("$H_2O$") are indicated with asterisks (*$P < 0.05$).

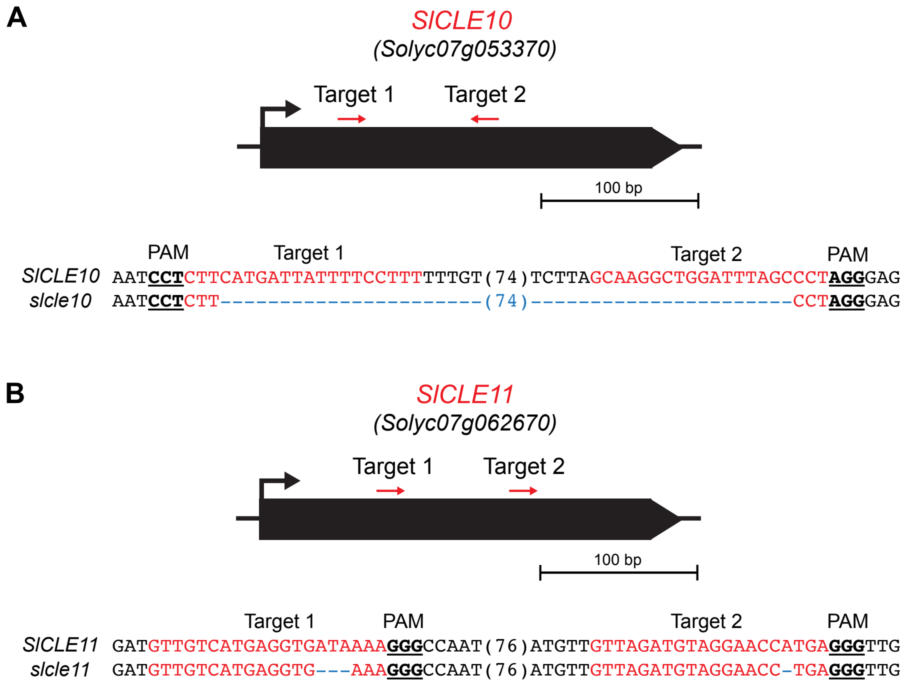


**Suppl figure 2. Outline of CRISPR guide RNA and mutations** introduced in (A) *Slcle10* (large deletion and introduction of stop codon), (B) *Slcle11* lines (deletion of isoleucine (32) and introduction of stop codon)

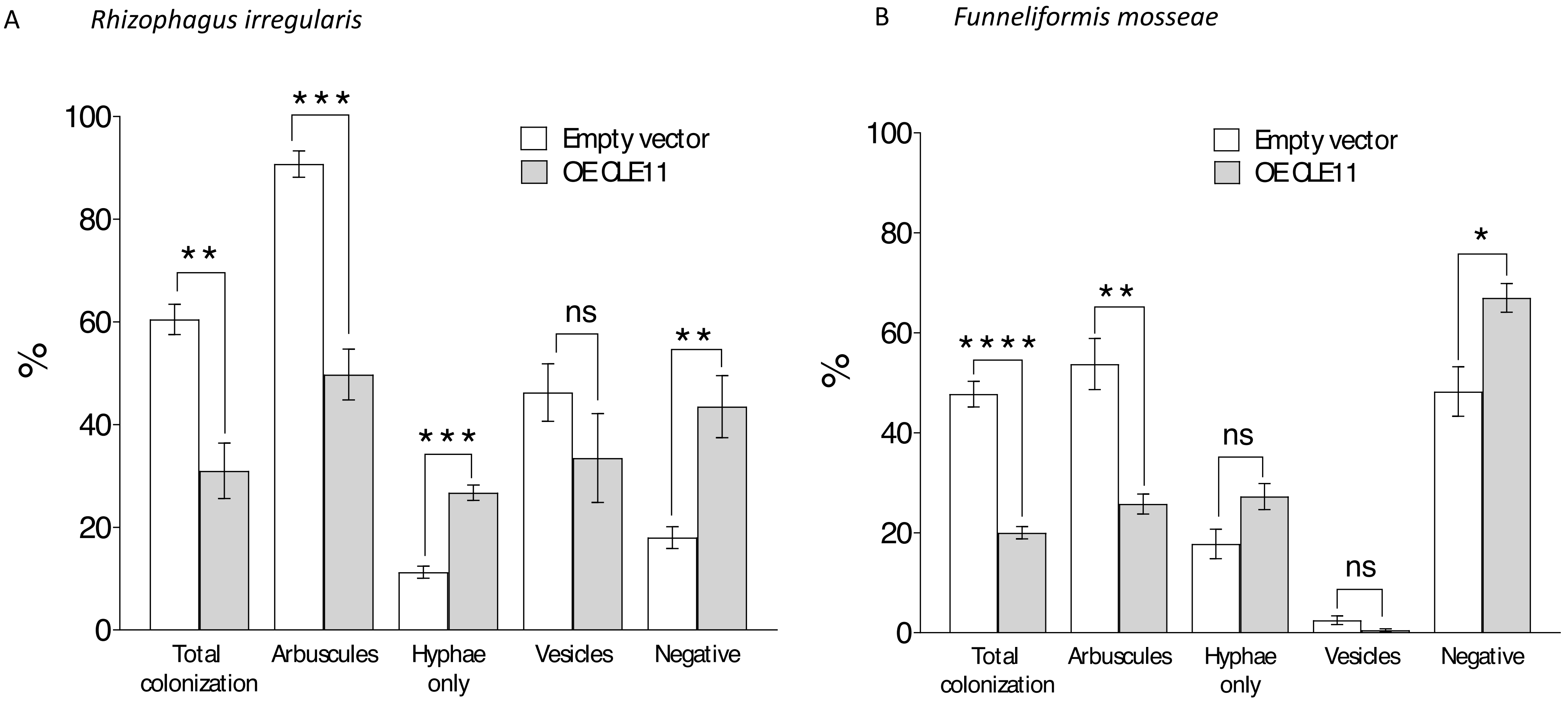


**Suppl. Fig 3. Mycorrhizal phenotype in *SlCLE11* overexpressing hairy roots.** *SlCLE11* OE and control hairy roots from composite tomato plants at 85 days after inoculation with the AM fungus *Rhizophagus irregularis* (A) or *Funneliformis mosseae* (B) were stained with trypan blue and the mycorrhizal phenotype was analysed. The percentage of root intersects without colonization ("negative"), just with hyphae ("hyphae only"), or with the presence of arbuscules and/or vesicles was calculated based on the McGonigle et al. (1990) procedure (n = 3; 120 root intersects per biological sample). Values correspond to mean ± SE. Significant differences (Student's t-test) between the *SlCLE11* OE-transformed plants and the control plants transformed with the corresponding empty vector are indicated with asterisks (**P* < 0.05; ***P* < 0.01; ****P* < 0.001; *****P* < 0.0001).
.

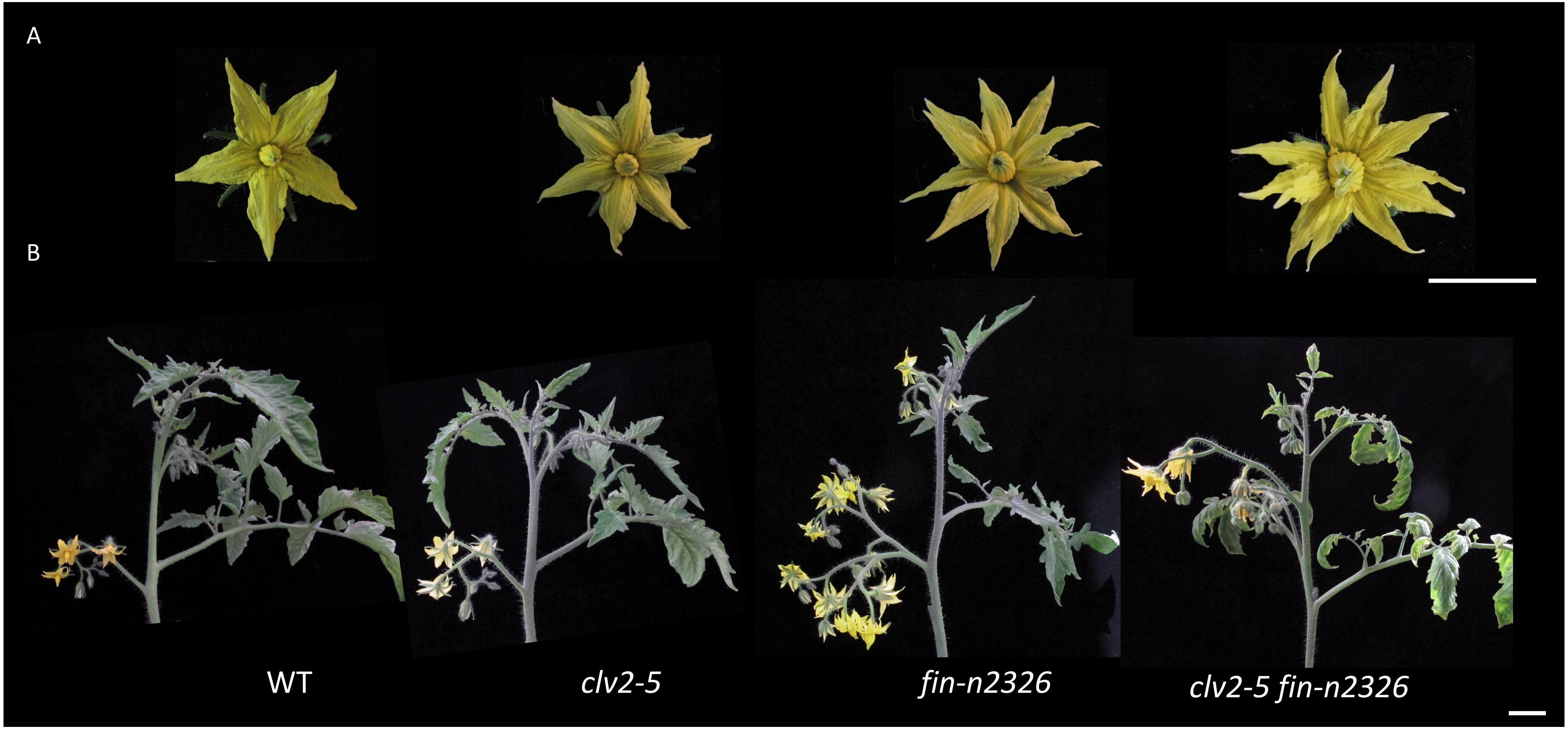


**Suppl Fig 4. Photos of shoot and flower phenotypes of WT (M82), *clv2, fin* and *clv2 fin* double mutant plants.** A. Individual flowers, B. inflorescence and apex. Scale bars are 1cm.

Suppl Table 1. The qRT-PCR and HRM primers used in the study. Some of the primer sequences are same as Zhang et al. (2014).

| Gene Name | Gene Locus | Primer | 5'-3' |
|---|---|---|---|
| *SlCLE1* | Solyc01g014100.2.1 | F | TCCAGATAAAGAGAATCAAGGAAGA |
| | | R | CGATTATGATGAAGCGGGTCT |
| *SlCLE2* | Solyc01g098890.1.1 | F | GAGGTCGCGGGTACCTAATG |
| | | R | AGGGATCAGGAGATGAGGGC |
| *SlCLE3* | Solyc02g067550.1.1 | F | GCATTCTCTTCAAAGGACCAATC |
| | | R | AGCTCGGATGCGTGTTATC |
| *SlCLE4* | Solyc02g087470.2.1 | F | GGCACTCCAAACAGGGAAGG |
| | | R | GGCATTTGGTCCAGTAGGCA |
| *SlCLE5* | Solyc03g025960.1.1 | F | AACCTCCCACTTCATTACTTCTTC |
| | | R | ATGATCTGCAGCACCAGCAT |
| *SlCLE6* | Solyc05g006610.2.1 | F | TGGAGGTGTTACAACAAAAATGA |
| | | R | GAACATGATGAGCACCACTTGA |
| *SlCLE7* | Solyc05g007650.1.1 | F | TGGTTTTCATTAGTAACAAGGCAT |
| | | R | ATACGCCTACTCGTCGATAACC |
| *SlCLE8* | Solyc05g053630.1.1 | F | CCCGTCTTGTCGGTTTTTCG |
| | | R | ACAGGATTTGCACCACTTGGA |
| *SlCLE10* | Solyc07g053370.1.1 | F | TGGAGGGTCGTATTCTTGATG |
| | | R | TTAGTCGGAGGCGAAGAGTG |
| *SlCLE10* genotyping | | F | TTAATCTTCACCTATGGCTAA |
| | | R | CTAGTTAGTCGGAGGCGAAGA |
| *SlCLE11* | Solyc07g062670.1.1 | F | ACGCGAATTTGGTTACGATGA |
| | | R | ATTGCGAGTGATGTTGAGCA |
| *SlCLE11* HRM | | F | CAAGGATGTTGTCATGAGGTG |
| | | R | CGTAACCAAATTCGCGTAAA |
| *SlCLE12* | Solyc09g061410.1.1 | F | TGATGGATATTGATCTCTTGTGGA |
| | | R | ATGAATGGTTGGGAACTGGAT |
| *SlCLE13* | Solyc09g091810.1.1 | F | ACGACCATGACCACGACTAT |
| | | R | ACTCCGGATTTGCACCACTA |
| *SlCLE14* | Solyc11g066120.1.1 | F | TTCATTCCCATGGCTCAACCT |
| | | R | GGATTCGGTCCAGATGGTGG |
| *SlCLE15* | Solyc11g071380.1.1 | F | GTGAAACTCCTAAACAGAAAGGTT |
| | | R | GAACTCCTCTTAGCTCCCAATC |
| *SlCAC* | Clathrin adaptor complexes | F | CCTCCGTTGTGATGTAACTGG |
| | medium subunit | R | ATTGGTGGAAAGTAACATCATCG |
| *SlEF1a* | Elongation factor 1 alpha | F | GCGTTGAGACTGGTGTGAT |
| | | R | GATGATGACCTGGGCAGTG |
| *SlFIN* genotyping | Solyc11g064850 | F | CAAACTTGGCTCGTGGAGAT |
| | | R | ATCATGGGTCTGCCACTAGC |
| *SlCLV2* HRM | Solyc04g056640 | F | TGATACTAATGCAACTGGGGTGT |
| | | R | CAAAAGCAAACCCATGTCCT |
